# Iterative, multimodal, and scalable single-cell profiling for discovery and characterization of signaling regulators

**DOI:** 10.1101/2025.08.27.672635

**Authors:** John D. Blair, Alexandra Bradu, Carol Dalgarno, Isabella N. Grabski, Rahul Satija

## Abstract

Cell signaling plays a critical role in regulating cellular state, yet uncovering regulators of signaling pathways and understanding their molecular consequences remains challenging. Here, we present an iterative experimental and computational framework to identify and characterize regulators of signaling proteins, using the mTOR marker phosphorylated RPS6 (pRPS6) as a case study. We present a customized workflow that uses the 10x Flex assay to jointly profile intracellular protein levels, transcriptomes, and CRISPR perturbations in single cells. We use this to generate a “glossary” dataset of paired protein–RNA measurements across targeted perturbations, which we leverage to train a predictive model of pRPS6 levels based solely on transcriptomic data. Applying this model to a genome-wide Perturb-seq dataset enables *in silico* screening for pRPS6 and nominates novel regulators of mTOR signaling. Experimental validation confirms these predictions and reveals mechanistic diversity among hits, including changes in signaling output driven by anabolic activity, cellular proliferation and multiple stress pathways. Our work demonstrates how integrated experimental and computational approaches provide a scalable framework for multimodal phenotyping and discovery.

## Introduction

Cell signaling orchestrates the critical decisions that underlie cellular identity, function, and fate. Rapid signaling dynamics frequently cascade into durable shifts in gene expression, linking pathway activity to both normal physiology and diseases such as cancer ^1^ and neurodevelopmental disorders ^2^. Yet the transcriptional effects of signaling are often subtle and co-occur with broader environmental and genetic factors, making it difficult to isolate direct causes from observational data ^3^. Discovering new regulators is equally challenging, and even when candidates are found, their functions may remain uncertain. To address these gaps, new approaches are needed to connect signaling perturbations with transcriptional responses at single-cell resolution, separate primary effects from secondary programs, and define the roles of uncharacterized regulators.

Pooled bulk CRISPR screens have proven to be powerful tools for discovering new regulators of signaling pathways ^4^. For example, genome-wide CRISPR knockout, interference or activation screens have identified key players in cell growth ^5^, DNA damage response ^6^, or immune activation ^7^ by linking perturbations to bulk phenotypic readouts. These approaches are most effective when leveraging essentiality phenotypes ^8^, cell surface markers ^9^, or reporter genes ^10^, but can struggle with more subtle or noisy phenotypes, particularly for intracellular markers such as phosphoproteins. In these cases, low signal-to-noise ratios and the technical challenge of preserving both high-quality antibody staining and genomic DNA for guide sequencing can limit sensitivity ^11^. As a result, bulk screens - especially when performed phenotypes based on intracellular protein staining - may yield incomplete or noisy lists of candidate regulators, obscure population heterogeneity, and fail to capture regulators active only in specific cell subsets, while also identifying candidates without molecularly characterizing their effects ^4,12^.

Advances in single-cell sequencing have enabled new strategies for multimodal analysis, including methods such as inCITE-seq ^13^, NEAT-seq ^14^, SIGNAL-seq ^15^, QuRIE-seq ^16^, and Perturb-icCITE-seq ^17^, which aim to capture cellular transcriptomes alongside intracellular or intranuclear proteins. While these approaches represent important progress, key limitations remain. First, these technologies rely on polyT-mediated reverse transcription performed after fixation and staining, a process that requires intact full-length RNA and can be highly variable across cell and tissue types, particularly in RNAse-rich contexts ^18^. Second, many are designed for nuclei rather than whole cells, which complicates perturbation experiments since capturable guide RNAs can be rapidly exported from the nucleus ^19^. These technical challenges reduce both gRNA recovery and transcriptome quality, underscoring the need for new methods that can robustly profile intracellular proteins in conjunction with perturbations at single-cell resolution.

Combining the rich multimodal data from single-cell sequencing with the scalability and unbiased scope of genome-wide bulk screens would enable this goal. Given the high dimensionality and information content of single-cell transcriptomics, we propose that computational modeling could help bridge this gap. Ideally, models could identify and predict signaling regulators from large-scale perturbation dictionaries, such as the growing compendium of genome-scale Perturb-seq datasets ^20,21^. This approach, however, would require the generation of new datasets that could link quantitative measurements of cell signaling activation with transcriptomic state at single-cell resolution.

Here, we present an integrated experimental and computational workflow to relate genetic perturbations with cell signaling level at genome-wide scale. We first introduce an experimental approach (FlexPlex) to perform multiplexed measurements across modalities, perturbations, and samples using the 10x Flex Single Cell assay. In parallel, we introduce a modeling approach (*‘in-silico’* Perturb-seq) that builds models of signaling output from scRNA-seq data and applies this to large Perturb-seq datasets. We utilize pooled genetic screens, FlexPlex, and *‘in-silico’* Perturb-seq to systematically identify regulators of mTOR signaling based on phosphorylated ribosomal protein S6 (pRPS6) levels. Our approach not only enumerates potential regulators but also provides both orthogonal validation and molecular insight into their function, enabling us to identify how the regulators either directly influence pRPS6, or indirectly influence signaling via growth or stress pathways. More generally, our approach demonstrates how multimodal single-cell technologies can be paired with computational modeling for efficient and large-scale identification and characterization of signaling regulators.

## Results

### FlexPlex quantifies protein, transcripts and perturbations in single cells

We aimed to develop a single-cell profiling workflow tailored for performing screens on intracellular phenotypes. To address key limitations with existing technology, in particular a reliance on reverse transcription and a focus on intranuclear instead of intracellular profiling, we utilized the 10x Genomics Flex assay. This workflow uses a transcriptome-wide probe set capable of capturing both full-length and degraded RNA and is inherently compatible with fixation required for intracellular staining. While the Flex assay is primarily used for the generation of gene expression data, custom extensions allow for quantifying additional features either by adding custom probes or by utilizing DNA oligos, such as ‘cell hashtag oligos’ (HTO) or ‘antibody-derived tags’ (ADT) with a pre-established capture sequence.

We leverage each of these custom modifications in our workflow which we refer to as ‘FlexPlex’ (full description in Supplementary Methods). Here, we introduce and optimize a ‘pre-fixation’ step, in which cells are first gently fixed and permeabilized, then incubated with antibodies targeting intracellular, intranuclear, and surface proteins (Fig. 1A). We subsequently fix again and incubate the sample with two types of RNA probes, the manufacturer-provided transcriptome-wide probe set and custom-designed probes targeting the highly expressed PolIII-gRNA transcripts. For intracellular protein detection, we and others ^22–24^ have previously developed custom conjugation strategies using DNA-tagged antibodies, which we show is compatible with the Flex assay. Immediately prior to antibody staining, we can also introduce unmodified single-stranded DNA oligos, or hashtag oligos (HTOs), which label distinct pools of cells and facilitate sample multiplexing ^25,26^. The resulting workflow is compatible with widely available commercial reagents and has been optimized to sensitively and simultaneously capture diverse molecular modalities which can then be selectively amplified using modality specific PCR handles (Table S1).

**Figure 1:**
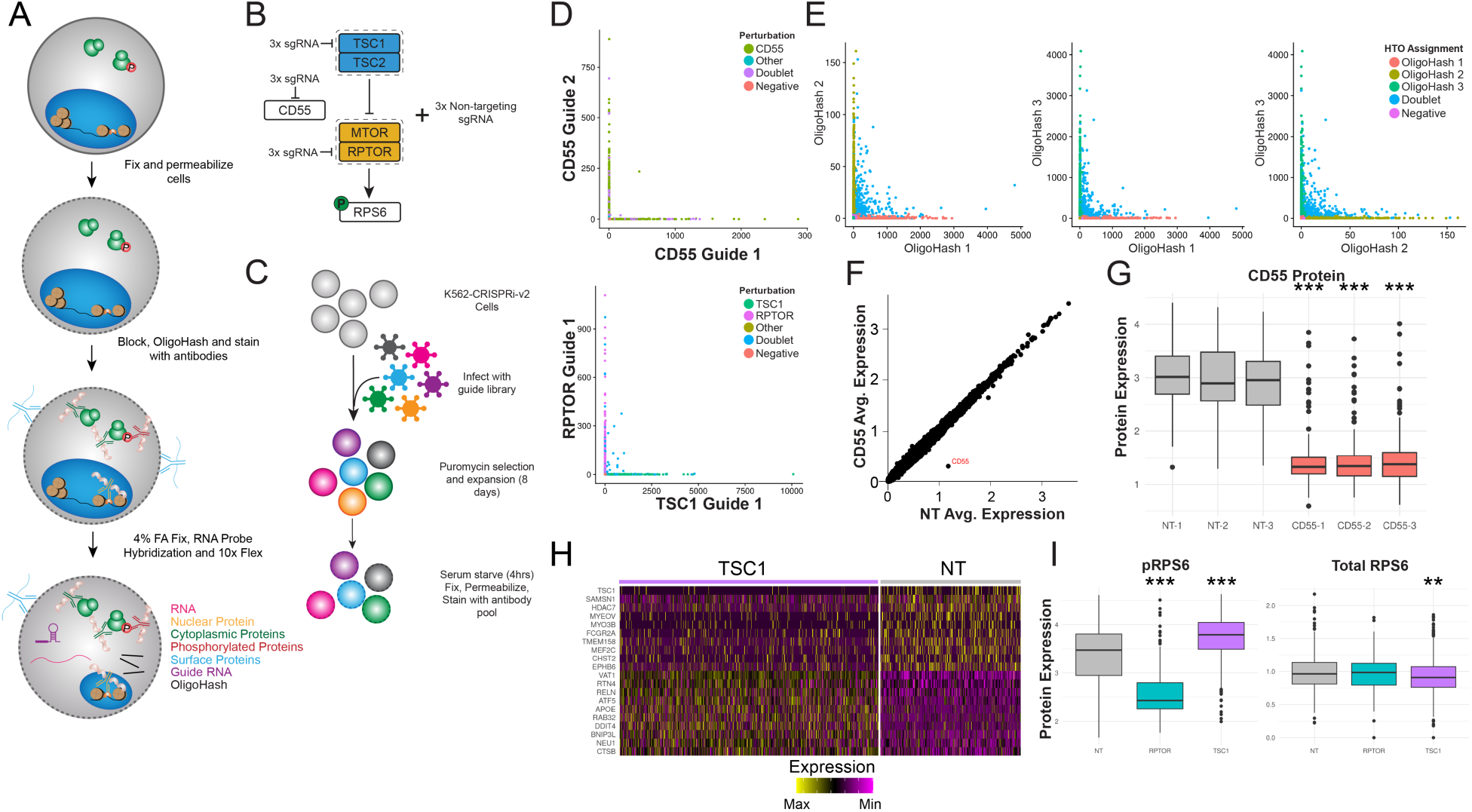
**a)** Schematic of FlexPlex experimental procedure. **b)** Experimental paradigm for FlexPlex pilot. **c)** Simplified schematic of perturbation strategy for pilot experiment. **d-e)** Scatter plots showing mutually exclusive expression of gRNA (d) and HTO (e) **f)** Scatter plot of average gene expression for each gene between non-targeting cells and CD55 perturbed cells. **g)** Boxplot of CD55 protein quantification between non-targeting and CD55 perturbed cells. **h)** Heatmap of up and downregulated genes between TSC1 perturbed cells and non-targeting cells. **i)** Boxplots of protein levels for phosphorylated RPS6 and total RPS6 in non-targeting, RPTOR and TSC1 perturbations. All statistical tests are two-sided t-tests adjusted using Benjamini-Hochberg correction for multiple comparisons. Significant differences between perturbations and non-targeting controls indicated by *** p <0.001,**p<0.01,*p<0.05. For all boxplots, the upper and lower bounds represent 75th and 25th percentiles respectively and the center line represents the median.

To evaluate its performance, we designed a pilot experiment to simultaneously profile RNA, intracellular proteins, cell surface proteins, guide RNAs (gRNAs), and HTOs ^25,26^. We applied FlexPlex to K562 cells engineered to express CRISPRi machinery and transduced with gRNAs targeting two key regulators of mTOR signaling ^27^,*TSC1* and *RPTOR*, as well as the surface protein *CD55* (Fig. 1B, C). The use of cell hashing substantially reduces per-cell library costs as it enables overloading of the 10x Chromium system followed by sample demultiplexing and doublet filtration (Supplementary Methods)^25^.

Across this experiment, we obtained high-quality data for all four modalities. UMI counts averaged approximately 7,000 per cell for RNA, 2,000 per cell for protein, 300 per cell for gRNA, and 200 per cell for cell hashing HTOs (Fig. S1A). Captured gRNAs were mutually exclusive within individual cells (Fig. 1D), enabling unambiguous perturbation assignment, while at the same time, hashing data facilitated reliable identification and removal of doublets (Fig. 1E). The resulting multimodal dataset clearly resolved the biological effects of each perturbation: *CD55* disruption led to highly specific and reproducible depletion of both CD55 RNA and protein (Fig. 1F,G), while perturbation of *TSC1* and *RPTOR* induced transcriptome-wide gene expression changes as well as alterations in pRPS6 levels. As expected, these changes could not be explained by measured differences in total RPS6 levels, validating our intracellular protein measurements (Fig. 1H,I).

### Genome-wide pooled screens identify regulators of RPS6 phosphorylation

Discovering novel regulators of phenotypes of interest requires an unbiased, genome-wide perturbation approach. While this approach is possible for FlexPlex, the practical requirements— processing multiple millions of cells and synthesizing a custom probe set for over 50,000 gRNAs—are likely to be cost-prohibitive. We therefore propose an alternative approach iterating between experiments and computational modeling that is cost-efficient and can be routinely applied (Fig. 2A).

**Figure 2:**
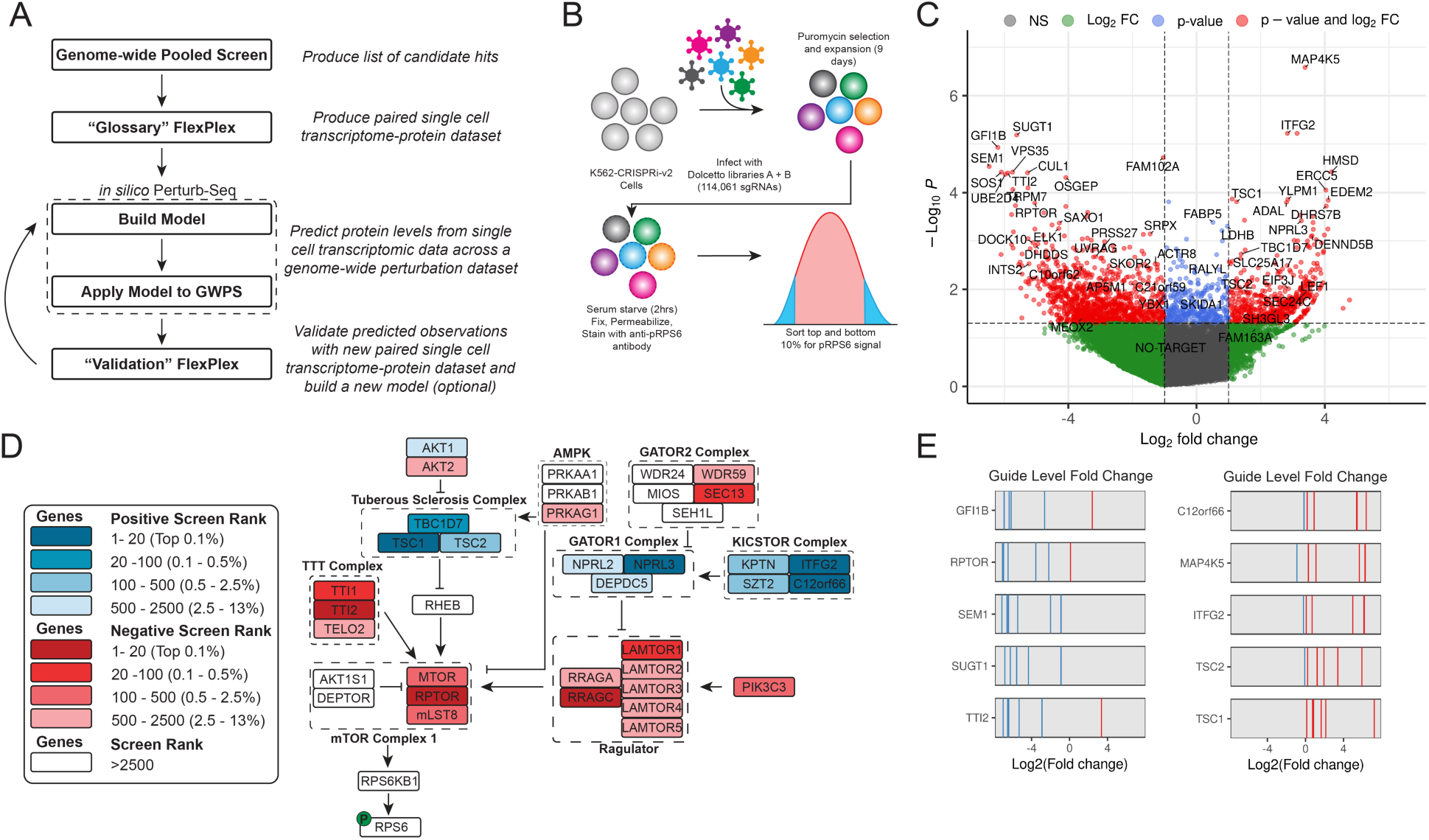
**a**) Iterative experimental–computational workflow: a genome-wide pooled CRISPR screen produces candidate hits, followed by a focused “glossary” FlexPlex experiment to pair protein and transcriptome data, which is then used to build predictive models. These models are applied to genome-wide Perturb-seq (GWPS) datasets and validated with additional FlexPlex experiments. **b)** Experimental design for the genome-wide pooled CRISPRi screen measuring intracellular phosphorylated RPS6 (pRPS6). **c)** Volcano plot showing fold-change and adjusted p-value for all perturbations, comparing high versus low pRPS6 fractions. **d)** mTOR pathway diagram with components colored by their pooled screen enrichment score (RRA rank) in the high or low pRPS6 fractions. **e)** Guide-level fold changes for selected hits, with red lines indicating positive fold changes (enriched in high-pRPS6 fraction) and blue lines indicating negative fold changes (enriched in low-pRPS6 fraction).

This multi-step strategy begins with a genome-wide pooled CRISPRi screen to generate an initial, although potentially noisy, list of candidate regulators. These candidates are then tested in a follow-up FlexPlex experiment, which pairs each cell’s transcriptome with protein measurements to create a “glossary” of protein–transcript relationships. This dataset trains a model that predicts protein expression from transcriptomic data alone, enabling large-scale *in-silico* screening of single-cell perturbation datasets to infer protein-level changes and nominate new candidates. These can be validated in subsequent FlexPlex experiments, iteratively expanding and refining the model (Fig. 2A).

We implemented this approach using intracellular phosphorylated ribosomal protein S6 (pRPS6) levels as a cellular phenotype. Biologically, pRPS6 is a downstream effector of mTOR signaling, which integrates a wide array of nutrient and growth factor cues, making it a sensitive readout of overall cellular activity ^27^. Technically, pRPS6 is a well-characterized post-translational modification with high-quality antibodies that have been validated in both bulk CRISPR screens^28^ and single-cell assays ^23,29^. Moreover, the mTOR pathway includes numerous well-defined regulators (e.g., *TSC1*, *RPTOR*), providing internal benchmarks for screen performance. Using CRISPRi K562 cells, we performed a genome-wide screen (∼19,000 genes; 114,061 gRNAs), fixed and stained cells for pRPS6, and FACS-sorted the top and bottom 10% to identify positive and negative regulators (Fig. 2B;S2A). Effects were quantified using Robust Rank Aggregation scores (Table S2) ^30^, fold-change, and statistical significance (Fig 2C; Supplementary Methods).

As expected, known repressors of mTOR signaling, including the Tuberous Sclerosis Complex (TSC1/2), and GATOR1 complexes were enriched in the high-pRPS6 fraction, reflecting increased mTOR activity upon their knockdown. Conversely, positive regulators such as components of mTORC1 (e.g., RPTOR, MLST8), the Ragulator complex (LAMTOR1–5), and the GATOR2 complex were enriched in the low-pRPS6 fraction, consistent with decreased signaling when these components are disrupted (Fig. 2C-E; Table S2). When comparing these results to a curated mTOR pathway diagram, we observed strong concordance across the pathway: nearly every major complex showed directional agreement between known biological function and screen enrichment (Fig 2D) ^27^. This included not only canonical core components of mTOR signaling, but also regulators of nutrient sensing (e.g., KICSTOR, AMPK) and mTORC1 complex assembly (e.g. TTI complex). This broad agreement reinforces the biological specificity and sensitivity of our intracellular screening platform for recovering both positive and negative regulators of a phospho-protein phenotype, even in the context of a complex signaling network.

In addition to positive controls, several of our hits (Fig. 2C) confirmed limited prior evidence linking them to mTORC1 function. For instance, TTI2, a component of the TTI1–TTI2–TELO2 complex, is known to stabilize and assemble members of the mTORC1 complex, including mTOR and RPTOR ^31^. Another hit, MAP4K5, is part of the MAP kinase pathway and has been implicated as an upstream modulator of mTOR signaling ^32^. More strikingly, we also recovered hits with no previously reported connection to mTORC1 activity. For example, GFI1B, a transcription factor involved in hematopoietic lineage differentiation ^33^, and DENND5B, a protein involved in vesicular transport and lipid metabolism ^34^, were both significantly enriched. The functional diversity of these hits, from transcriptional regulation to intracellular trafficking, suggests that RPS6 phosphorylation is influenced by a wide range of cellular processes.

Although our screen successfully identified numerous positive controls, some mTOR pathway members such as RHEB were missing, and others, like TSC2, ranked lower than anticipated given their known impact on mTOR signaling (Fig. 2C). Moreover, while top hits replicated in multiple independent experiments (Fig. S2B), genome-wide reproducibility across replicates was limited (R=0.20), but was consistent with the genome-wide reproducibility of a previously established pRPS6 bulk screen in HEK293 cells (R=0.25) ^28^. These discrepancies likely reflect common sources of noise in pooled antibody-based screens ^4^, including heterogeneity in staining across single cells, guide dropout, and uneven PCR amplification and highlight the need for additional tools to validate and refine these findings.

### Predicting pRPS6 levels from transcriptomic data

Building on our genome-wide screen results, we sought to determine if the heterogeneity in pRPS6 regulators could be captured and contextualized by our FlexPlex workflow. To test this, we performed a FlexPlex experiment in CRISPRi-v2 K562 cells transduced with a focused gRNA library targeting 50 genes (two gRNAs per gene) along with ten non-targeting controls (Fig. 3A). Targets were chosen from the pooled screen and aimed to span a spectrum of pRPS6 phenotypes, from high-confidence hits to lower-confidence candidates. We captured 4,577 gRNA singlets and assessed each perturbation’s effect on the transcriptome and pRPS6 levels. This dataset recapitulated a broad dynamic range of pRPS6 levels across perturbations (Fig. 3B) and between guides (Fig. S3A-C), consistent with expectations from the pooled screen. Interestingly, we found significant diversity in the transcriptional responses of different regulators, even those that induced similar changes in pRPS6 levels. Looking at the top differentially expressed genes induced by each perturbation (Fig. 3C,D) revealed strong similarity between TSC1- and TSC2-perturbed cells, consistent with their shared role in the Tuberous Sclerosis Complex but more broadly, most perturbations displayed distinct transcriptional signatures, even when their effects on pRPS6 were similar, underscoring the molecular heterogeneity of pRPS6 regulation.

**Figure 3:**
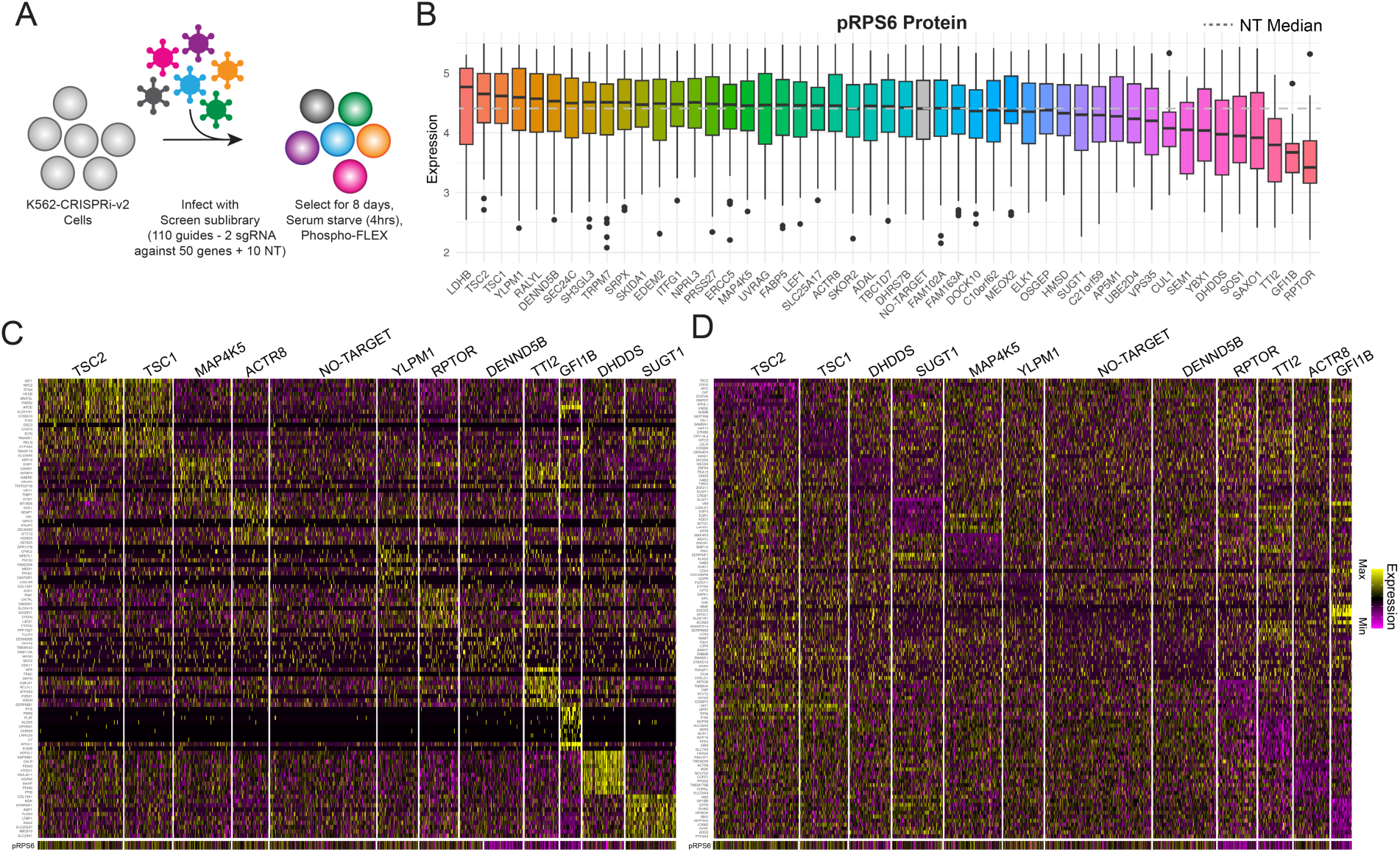
**a)** Experimental setup for “glossary” experiment. **b)** Boxplot of measured pRPS6 ADT values across all perturbations. Upper and lower bounds represent 75th and 25th percentiles respectively and the center line represents the median. **c)** Heatmap of upregulated genes and pRPS6 for representative perturbations in the “glossary” dataset. **d)** Heatmap of downregulated genes and pRPS6 for representative perturbations in the “glossary” dataset.

We reasoned that by pairing protein measurements with scRNA-seq across a diverse set of perturbations, this dataset could serve as a ’glossary’ (Fig. 2A) to link gene expression with pRPS6 levels. We therefore aimed to train a model to predict pRPS6 levels from transcriptomic data, which could serve as a surrogate readout of mTOR activity in contexts where direct protein measurements are unavailable. Our primary goal was to train a predictor in K562 cells that could be applied to existing genome-wide Perturb-seq (GWPS) datasets to systematically infer the effects of thousands of perturbations on mTOR signaling. However, given the central role of mTOR in cell biology, we also aimed to assess whether such a model could generalize to additional cellular contexts.

We explored several linear modeling strategies to build a reliable predictor while minimizing the possibility of overfitting and systematically evaluated their performance. We began with an ordinary least squares model trained on the 1,000 transcripts most positively and negatively correlated with pRPS6 levels (Fig. 4A). We then implemented two regularized approaches commonly used to improve generalization: LASSO regression, which removes non-informative features by shrinking their coefficients to zero ^35^, and Ridge regression, which is an optimal modeling choice when many features contribute small effects to the output variable ^36^. In both cases we set hyper parameters via cross-validation (Supplementary Methods). We found that the Ridge model performed best when evaluated against measured pRPS6 levels in an independent held-out FlexPlex validation experiment (R=0.77; Fig. 4A; S4A), indicating strong predictive performance on new experiments.

**Figure 4:**
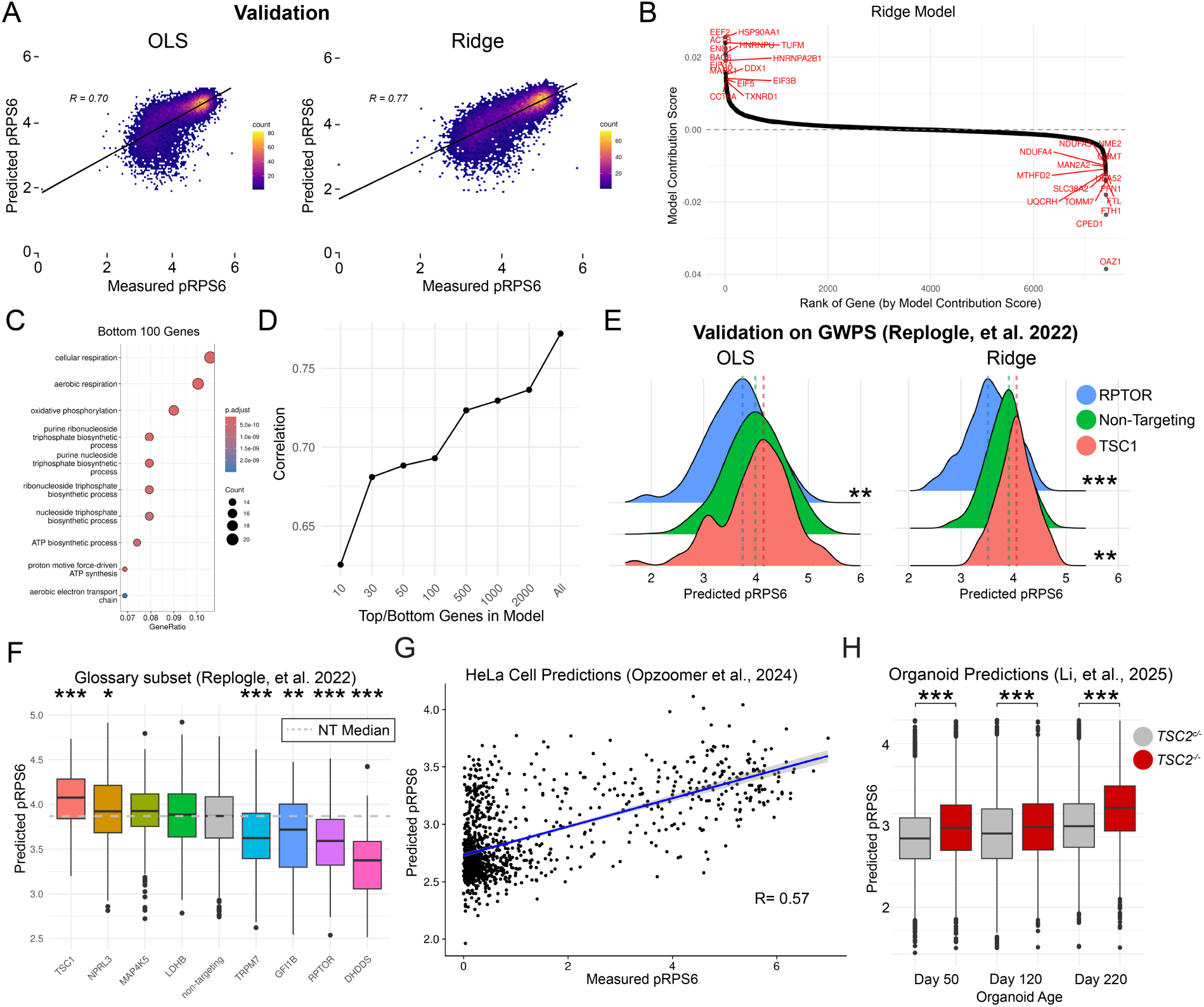
**a)** Correlation density plots comparing predicted pRPS6 to measured pRPS6 on a validation dataset across two linear modeling methods (OLS: R= 0.70; p < 0.0001; Ridge: R= 0.77; p< 0.0001). **b)** Rank-correlation plot of the model contribution score for all genes included in the Ridge pRPS6 model. **c)** Enriched gene ontology categories for top 100 positive regulators of pRPS6 based on model contribution score. **d)** Increasing the number of genes included in the Ridge model leads to an increase in the correlation between predicted pRPS6 to measured pRPS6 in a validation dataset. **e)** Density plots demonstrating the predicted pRPS6 differences between RPTOR, TSC1 and non-targeting perturbations across two linear modeling methods using transcriptional data from genome-wide perturb-seq (GWPS). **f)** Boxplot of predicted pRPS6 based on transcriptomic data from GWPS across a subset of perturbations present in the “glossary” experiment **g)** Scatter plot of predicted pRPS6 vs. measured pRPS6 in a HeLa cell dataset using SIGNAL-seq (ref 15) (R=0.57;p <0.0001). **h)** Violin plots of predicted pRPS6 levels amongst homozygous *TSC2*-knockout cells (TSC2^-/-^) versus heterozygous TSC2-knockout cells (TSC2^c/-^) in a separate brain organoid dataset from ref 37. All statistical tests are two-sided t-tests adjusted using Benjamini-Hochberg correction for multiple comparisons. Significant differences between compared to non-targeting controls indicated by *** p <0.001,**p<0.01,*p<0.05. For all boxplots, the upper and lower bounds represent 75th and 25th percentiles respectively and the center line represents the median.

We found that the 100 genes with the most negative model contribution scores (Supplementary Methods) were highly enriched for pathways related to cellular respiration and growth, consistent with the known role of mTOR in regulating these processes ^27^ (Fig. 4B,C). However, model performance continued to improve as additional genes were included, indicating that predictive accuracy was not solely driven by a small subset of core regulators (Fig. 4D). Notably, even among the top-weighted genes, associations were often context-dependent and occasionally counterintuitive. For example, lower expression of the elongation factor EEF2 correlated with reduced pRPS6 levels in K562 cells, yet this pattern was not universal as EEF2 was also downregulated in TSC1-deficient cells where we observed an increase in pRPS6 levels. These findings suggest that while transcriptomic models can robustly infer mTOR activity, their strength lies in leveraging distributed signals across the transcriptome rather than relying on a conserved core module of a few dominant genes.

Next, we tested the ability of our model to predict pRPS6 levels in externally generated experimental datasets and new biological contexts. First, we applied it to curated subsets of GWPS in K562 cells for both the mTOR pathway and the glossary perturbations. We observed significant predicted pRPS6 differences between different mTOR pathway perturbations, such as *RPTOR* and *TSC1*, as expected (Fig. 4E,S4B,C). Extending this validation analysis to all perturbations in the glossary experiment also revealed a strong correlation between GWPS-predicted and FlexPlex-observed pRPS6 levels (R=0.59; Fig. 4F;S4D).

Following this, we tested the model in a new biological context by applying it to an independent HeLa cell dataset in which both RNA and pRPS6 levels were measured at single-cell resolution^15^. Despite being trained exclusively on K562 data, our model achieved a single-cell correlation of R=0.57 between predicted and measured pRPS6 levels (Fig. 4G). Finally, to evaluate the generalizability beyond cancer cell lines, we applied it to a dataset of cortical spheroids derived from conditional TSC2-knockout iPSCs, sampled across multiple developmental timepoints ^2,37^. The model predicted elevated pRPS6 levels in homozygous TSC2 knockout spheroids at later timepoints (Fig. 4H), which had been independently validated by both immunofluorescence staining and western blot analysis and is consistent with the expected hyperactivation of mTOR signaling ^2,37^. We conclude that this modeling approach learns relationships between mTOR activity and transcriptomic state that can be extended beyond a single experiment or biological context.

### Predicting novel mediators of pRPS6 in single-cell genome wide perturb-seq

Having validated the pRPS6 prediction model across multiple experimental contexts, we next applied it to the full GWPS dataset to identify novel regulators of mTOR signaling. This application enabled us to expand phenotypic characterization across all 9,867 perturbations in the dataset, using predicted pRPS6 levels to prioritize candidate regulators for follow-up. We refer to this process, which was inspired by the original GWPS manuscript’s efforts to screen their dataset for inferred phenotypes extending beyond gene expression ^20^, as ’*in-silico*’ Perturb-seq (Fig. 5A).

**Figure 5:**
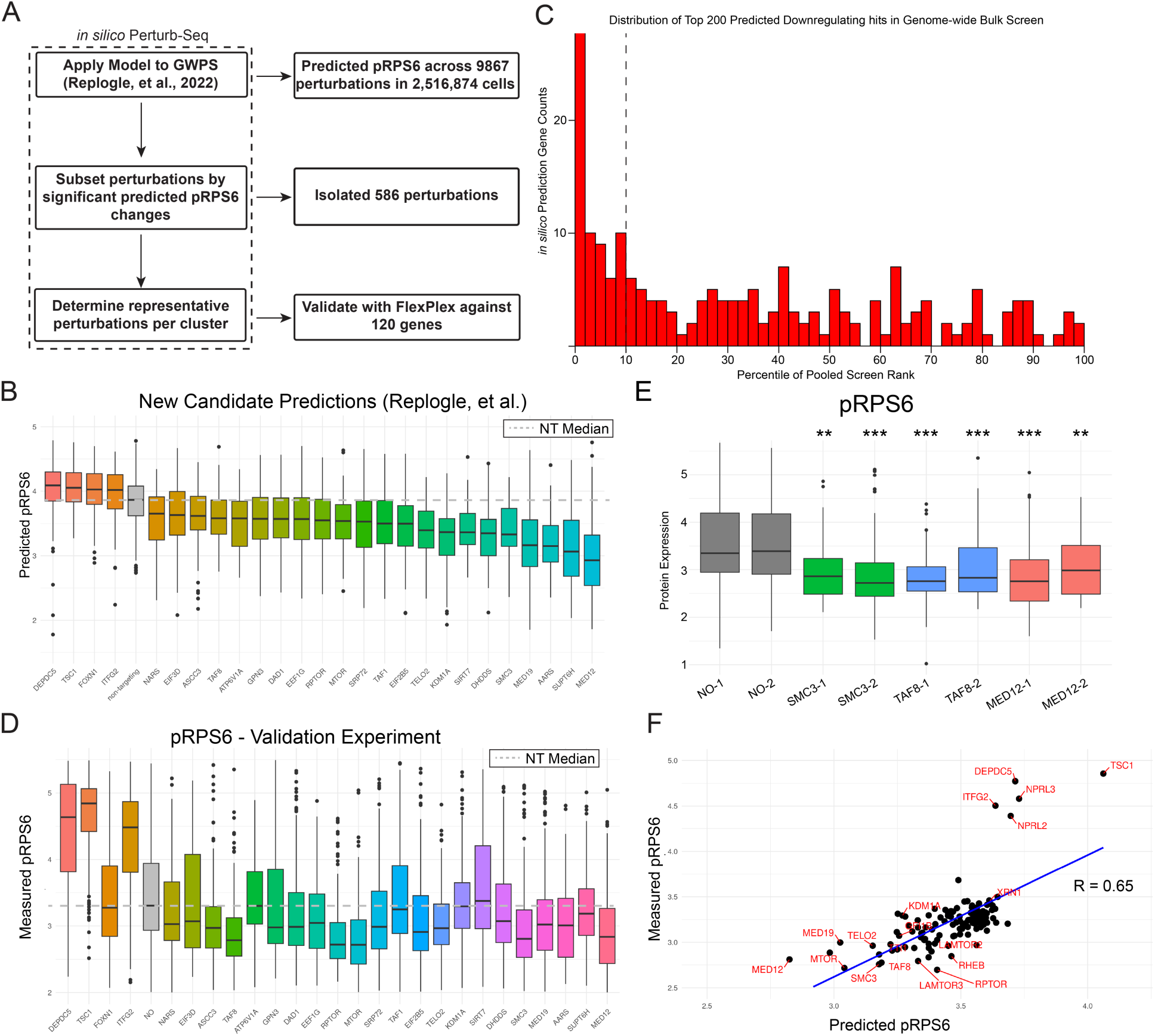
**a)** Schematic of the *in silico* Perturb-seq workflow. Predictions from genome-wide Perturb-seq (GWPS) were used to infer pRPS6 levels, select top candidate regulators, and validate them with FlexPlex. **b)** Predicted pRPS6 levels across a subset of GWPS perturbations**. c)** Distribution of pooled screen rank-percentiles for the top 200 predicted negative pRPS6 regulations, showing enrichment among high-ranking hits (Chi-Square test; p <0.001). **d)** Measured pRPS6 levels for the same perturbations in the FlexPlex validation experiment, plotted as in (b). **e)** Validation of reproducibility across independent gRNAs for selected regulators from the FlexPlex dataset. Protein expression values for pRPS6 are shown, with significance compared to non-targeting controls. **f)** Correlation of the medians of predicted pRPS6 and measured pRPS6 for each perturbation from the validation FlexPlex experiment (R = 0.65; p = 1.65 x 10^-15^). All statistical tests are two-sided t-tests adjusted using Benjamini-Hochberg correction for multiple comparisons. Significant differences between compared to non-targeting controls indicated by *** p <0.001,**p<0.01,*p<0.05. For all boxplots, the upper and lower bounds represent 75th and 25th percentiles respectively and the center line represents the median.

We identified a total of 586 perturbations predicted to significantly regulate pRPS6 levels (FDR<1%; Table S3; Fig. 5B). The identification of putative regulators skewed towards positive regulators (i.e. perturbations that resulted in a decrease of pRPS6), reflecting the serum-rich environment and correspondingly high baseline mTOR activity in which the GWPS K562 cells were cultured. Identified perturbations included known mTOR regulators (i.e. members of the GATOR1 and KICSTOR complexes) and were also highly enriched (p < 0.0001) for genes that fell within the top 10% of enriched regulators in our pooled pRPS6 screen (Fig. 5C). However, many of the identified regulators showed no enrichment in the pooled screen (Fig. 5C), suggesting that *in-silico* Perturb-seq could help to prioritize potential false negatives for additional experimental validation.

As expected, hits in our in-silico screen represented a diverse set of regulators and pathways (Table S3; Fig. S5A). For instance, we predicted depletion of pRPS6 levels upon perturbation of multiple members of the mediator complex. This result was intriguing because the mediator complex is involved in transcriptional regulation and has only loosely been linked to mTOR signaling ^38^, and no mediator complex members were included in our ’glossary’ experiment. Other identified factors included TAF8 and SMC3 which, like the mediator complex, generally regulate core cellular processes ^39,40^ but have not been specifically linked to mTOR. Additional hits included translation initiation factors (i.e. EIFs) and several aminoacyl-tRNA synthetases (aaARS genes), highlighting a broad translation-related program converging on pRPS6.

While these results suggest that *in-silico* Perturb-seq may be able to identify regulators of intracellular signaling programs that are missed by other approaches, our findings are ultimately computational predictions that require experimental validation. To address this, we selected a subset of 120 regulators spanning a range of predicted effects on pRPS6 signaling for follow-up. These regulators were chosen based on transcriptomic clustering of perturbations in the GWPS dataset followed by sampling across these clusters, allowing us to more evenly assess regulators in different transcriptional programs and maximize pathway diversity in the validation set (Supplementary Methods).

To empirically test our predictions, we performed an additional FlexPlex experiment for 250 gRNA targeting these 120 genes in 23,704 cells. This allowed for a direct comparison between the predicted effects from transcriptomic data and measured differences in pRPS6 levels between NT and perturbed cells. We observed clear validation of model predictions for both known controls and novel regulators (Fig. 5D). For instance, perturbation of MED12 led to a 63.4% reduction in pRPS6 levels (p = 2.37 x 10^-16^), consistently observed across independent gRNAs (Fig. 5E). Similar decreases were observed for SMC3 (33.6% reduction; p = 0.03) and TAF8 (53.1% reduction; p = 2.86 x 10^-9^), confirming these novel predictions and reinforcing the utility of *in-silico* Perturb-seq in identifying previously uncharacterized mTOR regulators.

Looking globally across all 120 tested regulators, we observed a strong positive correlation between measured and predicted pRPS6 levels (R = 0.65; p < 0.0001; Fig. 5F), supporting the predictive accuracy of the model. Notably, this validation was achieved despite experimental differences between the GWPS and validation datasets, including nutrient starvation and harvesting conditions. While these conditions resulted in shifts in the absolute magnitude of pRPS6 modulation, the ranking of perturbations was consistent: both the model and validation experiment identified perturbation of canonical negative regulators such as TSC1 and DEPDC5 as among the strongest inducers of pRPS6. Together, these results demonstrate that transcriptome-based modeling can effectively prioritize candidate regulators of mTOR signaling, even under varying experimental contexts.

FlexPlex can simultaneously measure multiple intracellular proteins, enabling parallel screens within the same experiment. While our study focused on pRPS6, which guided our initial set of perturbations, we also included Vimentin, an intermediate filament protein ^41^, in the glossary experiment. This allowed us to repeat the process of constructing a ridge regression model, predicting transcriptome-wide effects of perturbations from the GWPS dataset, and confirming these predictions in a FlexPlex validation experiment. Model accuracy, assessed at both the single-cell (R = 0.73; p < 0.0001; Fig. S5B) and perturbation (R = 0.73; p < 0.0001; Fig. S5C) levels, was similar to or exceeded what we observed for pRPS6. Interestingly, a subset of regulators of mTOR signaling also influenced Vimentin levels, though often in counterintuitive ways: for example, RPTOR and LAMTOR both decreased pRPS6 but had opposing effects on Vimentin, as did TSC1 and DEPDC5 (Fig. S5D). Our validation dataset confirmed multiple predicted regulators of Vimentin but as expected, did not identify VIM perturbation itself, since this perturbation alters only a single transcriptomic target (Fig. S5E). As a proof-of-concept, these results demonstrate how future studies can systematically profile multiple phenotypes in parallel.

### Deconvolving cellular process influencing pRPS6 levels

Building on our modeling results showing that distinct transcriptional programs can converge on the same phenotypic consequences for pRPS6, we sought to identify regulatory modules and pathways linked to mTOR signaling. We compiled differentially expressed genes from validated regulators, supplemented with curated markers of the integrated stress response (ISR) and unfolded protein response (UPR) from the literature ^42^. Performing k-means clustering of this combined set revealed seven transcriptional modules and their associated regulators (Fig. 6A).

**Figure 6:**
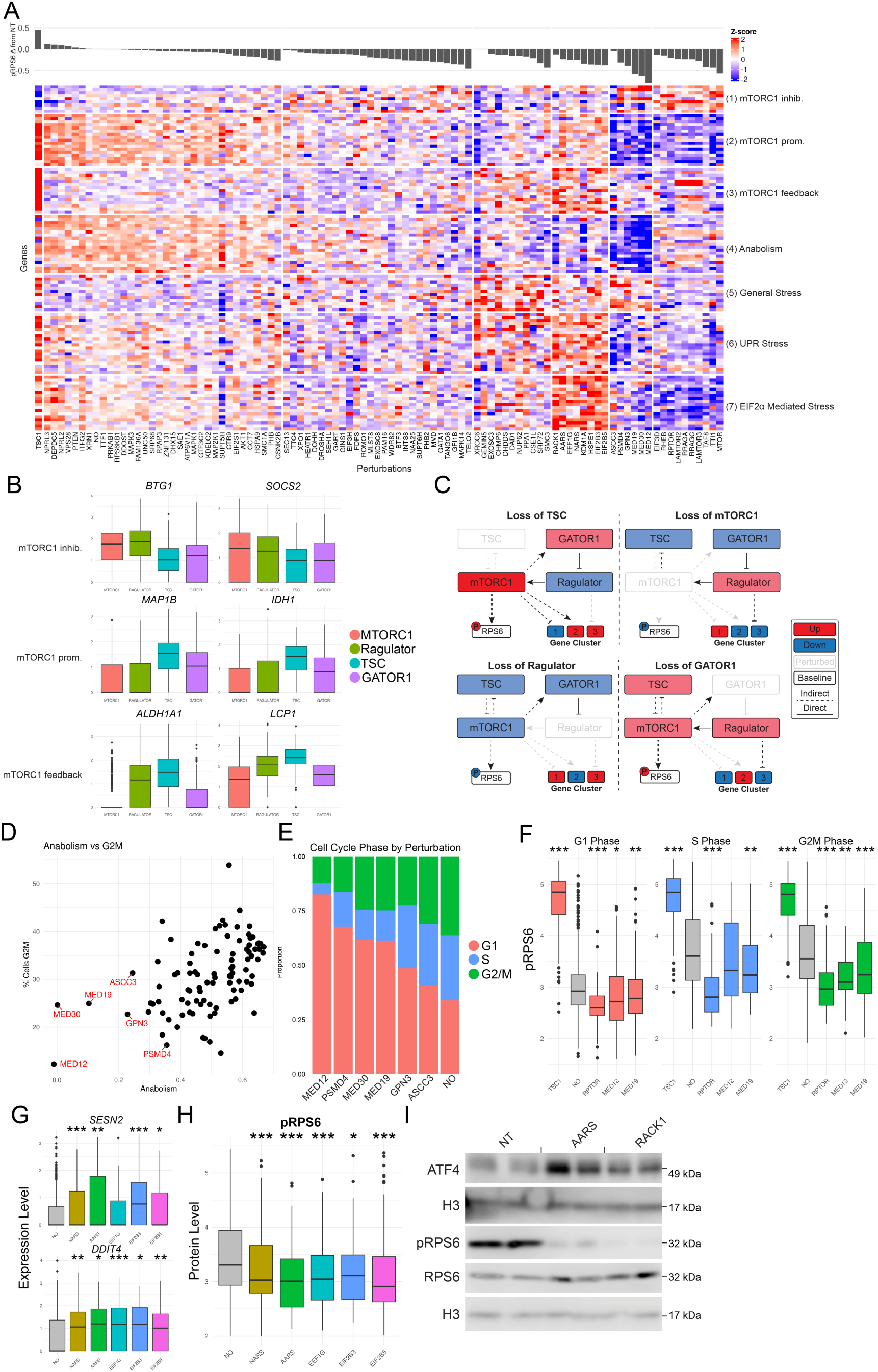
**a)** Heatmap of k-means–clustered gene modules across the validation dataset. Rows represent gene modules and columns represent perturbations, with the barplot above showing median predicted pRPS6 values per perturbation. **b)** Boxplots of representative differentially expressed genes for selected mTORC1 pathway perturbations, grouped by complex. **c)** Schematic models illustrating inferred feedback relationships among different mTORC1 pathway complexes. **d)** Scatter plot comparing median scores of the anabolism gene module with the fraction of cells in G2/M phase for each perturbation. **e)** Proportional bar plot of cell cycle phase distribution across perturbations. **f)** Boxplots of measured pRPS6 levels stratified by cell cycle phase for selected perturbations (MED12, MED19, TSC1, RPTOR) compared to non-targeting controls. **g)** Boxplots of ATF4-related transcripts across perturbations predicted to activate the ISR. **h)** Boxplots of measured pRPS6 levels across EIF2α-related perturbations. **i)** Western blot analysis of ATF4, total RPS6, and pRPS6 following individual perturbations in K562 cells. Representative data from two independent experiments are shown. All statistical tests are two-sided t-tests adjusted using Benjamini-Hochberg correction for multiple comparisons. Significant differences between compared to non-targeting controls indicated by *** p <0.001,**p<0.01,*p<0.05. For all boxplots, the upper and lower bounds represent 75th and 25th percentiles respectively and the center line represents the median.

Two gene modules (1 and 2) represented stereotypical patterns associated with the activation and inhibition of some key, but not all, regulators of mTOR signaling (Fig. 6A,B). For example, module 2 was sharply downregulated upon inhibition of positive mTOR regulators (i.e. MTOR and RAPTOR) but induced upon inhibition of TSC1. Consistent with our previous findings that no gene module could uniformly predict pRPS6 levels, this gene module was also counterintuitively up-regulated upon inhibiting translational regulators (i.e. EIF factors), which reduce pRPS6 levels. Genes in this module include MAP1B, a cytoskeleton regulator ^43^, and IDH1, a key enzyme for energy production ^44^, suggesting that this program reflects anabolic activity and biosynthetic demand. Module 1 exhibited opposite patterns, representing genes that were generally associated with low levels of mTOR signaling. Many of these genes, like BTG1 and SOCS2, help to inhibit cell growth and proliferation ^45,46^. A third gene module (module 3) showed upregulation in both TSC1 and Ragulator perturbed genes and downregulation in mTORC1 and GATOR1 perturbed genes (Fig. 6A,B), implying indirect transcriptomic feedback, and highlighting the complexities of mTORC1-related transcriptional regulation (Fig. 6C).

A separate transcriptomic gene module was enriched for genes involved in cellular growth and anabolic processes, including HSPs and EEF2, which support biosynthetic activity ^47,48^. This gene module was most strongly downregulated in cells subjected to perturbations that broadly affected transcriptional output, including mediator complex perturbations ^49^, and GPN3, which helps transport RNA PolII into the nucleus ^50^. While this gene module did not include cell cycle stage markers, we found that expression of this module across perturbations was correlated with cellular proliferation (Fig. 6D), and the percentage of actively cycling cells, as assessed by cell cycle stage analysis (Supplementary Methods; Fig. 6E).

More broadly, our single-cell measurements revealed that pRPS6 levels vary across cell cycle stages, providing an opportunity to disentangle whether observed reductions in pRPS6 arise from changes in cell cycle distribution or from direct effects on signaling within specific phases. Leveraging transcriptomic profiles to infer cell cycle stage (Fig. S6A,B), we examined pRPS6 levels within matched phases across different perturbations (Fig. 6F;S6C). This analysis revealed that, even after controlling for cell cycle stage, significant changes in pRPS6 levels persisted, not only for canonical mTORC1 regulators such as TSC1 and RPTOR, but also for members of the mediator complex. These findings indicate that observed reductions in pRPS6 reflect both an increased proportion of cells arrested in G1 and a true decrease in signaling levels within G1-phase cells themselves. We note that the ability to distinguish between these factors is specifically enabled by paired transcriptomic measurements, and that variation in cell cycle phase can be a confounding factor in bulk CRISPR screens.

In addition, we identified three gene modules associated with cellular stress responses. One module captured a general stress program marked by elevated expression of DDIT3 and PPP1R15A, particularly in perturbations that restricted cellular growth (Fig. S6D). A second module was enriched for genes activated by the ATF6 and IRE1α branches of the unfolded protein response (UPR) ^42^, highlighting both known UPR-related perturbations (e.g., SRP72, DHDDS, DAD1) and novel candidates (e.g., XRCC6, PPA1) (Fig. S6E). A third stress module reflected activation of EIF2α-mediated signaling, which can occur through the integrated stress response (ISR) or the PERK branch of the UPR ^42^. High expression of this module was observed in perturbations targeting core ISR regulators (EIF2B3, EIF2B5), as well as aminoacyl-tRNA synthetases (AARS, NARS), whose inhibition activates the GCN2 kinase via accumulation of uncharged tRNAs ^51^. EIF2α phosphorylation leads to selective translation of ATF4 and a downstream transcriptional cascade, including DDIT4 (an mTOR inhibitor) and SESN2 (involved in metabolic recovery) ^52^ (Fig. 6G). These perturbations consistently exhibited reduced pRPS6 levels (Fig. 6H).

We also identified additional perturbations with strong transcriptional similarity to canonical ISR regulators, but without prior links to the ATF4 pathway. These included EEF1G, a factor responsible for delivering aminoacyl-tRNAs to the ribosome ^53^, and RACK1, a component of the EIF3 complex ^54^. To test whether these candidates indeed induce ISR, we performed western blot analysis of ATF4 and pRPS6 following CRISPRi knockdown. We found that, like aminoacyl-tRNA synthetases such as AARS, knockdown of RACK1 led to robust induction of ATF4 and depletion of pRPS6. These results extend our transcriptomic observations, providing direct protein-level evidence that RACK1 perturbation can activate the ISR and further highlighting how previously uncharacterized regulators can converge on this pathway (Fig. 6I).

Together, these analyses deconvolve the layered contributions of cell cycle, anabolic growth, and stress response programs to intracellular pRPS6 levels. By integrating single-cell measurements of gene expression and signaling, we demonstrate that reduced pRPS6 can arise through diverse mechanisms including canonical mTORC1 inhibition, broad transcriptional repression, and activation of stress-induced translational control pathways. More generally, we demonstrate how multimodal single-cell data can be combined with genetic perturbation screens to identify novel regulators and disentangle their contribution to the modulation of intracellular signaling networks.

## Discussion

In this study, we introduce FlexPlex, a multimodal single-cell workflow that enables simultaneous profiling of transcripts, proteins, perturbations, and sample identity, and combine it with a computational framework, in-silico Perturb-seq, to identify regulators of intracellular signaling. Using pRPS6 and mTOR signaling as a case study, we demonstrate that transcriptomic information can be leveraged to predict protein levels and systematically identify and prioritize candidate regulators for validation.

A key aspect of our approach is the iterative cycle between experimentation and modeling. Rather than attempting a prohibitively expensive genome-wide multimodal screen, we combined complementary strategies: a pooled CRISPRi FACS screen to generate a broad but noisy list of candidate regulators, a targeted FlexPlex experiment to establish a high-resolution “glossary” linking transcriptomes with protein phenotypes, and computational modeling to extend those relationships in silico across genome-wide Perturb-seq datasets. This cycle can then be repeated, with FlexPlex validation refining the model and expanding its predictive scope. Such an iterative framework balances the breadth of pooled screening with the depth of single-cell multimodal profiling, providing a cost-efficient and scalable strategy for mapping intracellular signaling regulators.

Our results highlight that the relationship between transcriptome and protein phenotype is informative but not straightforward. Protein levels are not encoded by a simple one-to-one mapping from RNA expression but rather emerge from distributed signals across many genes. This was reflected in the superior performance of ridge regression, with optimal model performance occurring when integrating small contributions from thousands of features and was further supported biologically by our identification of multiple distinct pathways converging on pRPS6 regulation. These findings suggest that the full value of single-cell transcriptomics lies not only in identifying condensed signatures or small sets of differentially expressed genes, but also in leveraging the aggregate information embedded across the entire expression space.

These results underscore the potential of transcriptome-based prediction as a scalable framework. With the rapid expansion of multimodal single-cell sequencing techniques that include the transcriptome, the same approach can be readily applied to infer many different phenotypes and accelerate discovery across diverse biological processes. Indeed, we show in the same experiment that transcriptomic data can accurately predict levels of Vimentin, an intermediate filament protein, and leverage this to identify and characterize its regulators as well.

An important limitation is that our predictive models rely on perturbation dictionaries generated in the same cellular context, which constrains their immediate generalizability. We show that some cross-context applications are possible—for example, applying K562-trained models to HeLa cells or iPSC-derived spheroids—but predictive performance is inevitably reduced when models are transferred across biological systems. Expanding context-specific perturbation datasets will therefore be essential for broadening the scope and accuracy of this approach.

Encouragingly, there is significant emphasis on broadening these perturbation resources, particularly the development of large-scale Perturb-seq dictionaries across diverse cell types and conditions ^21,55,56^ as part of the field’s growing interest in constructing virtual cells ^57^, To date, these dictionaries have primarily focused on RNA readouts, but we propose that models trained on multimodal datasets can act as bridges, allowing the wealth of existing transcriptomic data to be reinterpreted in terms of cellular phenotypes that were not assayed in the original dictionary. In this way, single-cell atlases may serve not only as catalogs of gene expression, but also as platforms for virtual phenotyping, enabling systematic inference of protein activities, pathway states, and functional outputs across millions of cells.

## Methods

### Cell Culture

K562-CRISPRi-v2 cells, were derived as previously stated ^56^ and were maintained in IMDM media (ThermoFisher: 31980030) supplemented with 10% FBS (Corning:35-010-CV), 1 mM Pen-Strep (Sigma: P0781-50ML) and 1X Non-essential amino acids (Sigma:M7145-100ML). HEK293FT cells were acquired from Thermo Fisher (R70007) and maintained in DMEM media (Caisson Labs: DML10) supplemented with 10% FBS and 1X non-essential amino acids. All cells were grown at 37° C and 5% CO_2_.

### Guide RNA cloning

Human Dolcetto CRISPR Inhibition Pooled Library (Set A and B) was a gift from David Root and John Doench (Addgene #1000000114) ^58^. These libraries were expanded and purified as recommended.

For the pilot, glossary and validation experiments, guide sequences for genes of interest were selected from the full Dolcetto libraries and ordered from IDT as an oligo Pool (oPool) of top and bottom pairs with Esp3I integration sequences at either end (Top: CACCGNNNNNNNNNNNNNNNNNNNN, Bottom: AAACNNNNNNNNNNNNNNNNNNNNC.

Pooled guides were annealed and ligated into a Esp3I digested modified CROP-seq cloning vector ^19^ using standard guide RNA cloning practices as described ^59^. Ligated plasmids were then cleaned up and concentrated through isopropanol precipitation and resuspended in TE buffer. Concentrated plasmid libraries were then electroporated into Endura electrocompetent cells (Lucigen: 60242-1), recovered in warm SOC media for 1 hour and plated onto ampicillin-containing LB-Agar plates. The following morning, the lawn of bacterial colonies was harvested into Luria Broth and purified using a Midiprep kit (Zymo: D4200). For all libraries, guide RNA balance was verified through sequencing of the plasmid library on an Illumina MiSeq instrument.

### Lentiviral Production

For virus production, 10 million HEK293FT cells were seeded in a 10-cm dish 18 h before transfection. We used 45 µl 1 mg/ml polyethyleneimine (PEI; Polysciences 23966-1) and 20 µg total plasmid DNA (6.4 µg psPAX2: Addgene, #12260; 4.4 µg pMD2.G: Addgene, #12259; 9.2 µg sgRNA-expressing plasmid) per transfection reaction. Six hours post-transfection, the medium was exchanged for 10 ml DMEM + 10% FBS containing 1% bovine serum albumin (VWR 97061-420). Viral supernatants were collected after an additional 48 h, spun down to remove cellular debris for 5 min at 4 °C and 1,000 x g, passed through a 0.22 μm filter and stored at −80 °C until use.

### Antibody Conjugation

Antibody conjugation and pooling was undergone exactly as previously described ^23^. Briefly, single-stranded DNA oligos in the TotalSeq-B format (/5AmMC12/GTGACTGGAGTTCAGACGTGTGCTCTTCCGATCTNNNNNNNNNNXXXXXXXXX XXXXXXNNNNNNNNNGCTTTAAGGCCGGTCCTAGC*A*A) were ordered from IDT. We attached a TCO-PEG4 linker to each oligo through incubation with TCO-PEG4-NHS reagent (Click Chemistry Tools: A137-25) and then column purified them with Micro Bio-Spin 6 Columns (Bio-Rad: 732-6221).

Next, the purified antibody was labeled with mTz-PEG4 (Click Chemistry Tools: 1069-10) and conjugated to the TCO-PEG4 labeled oligo at a target concentration of 15 pmol per 1 μg of antibody. Finally, for each experiment, 3-5 μg of each antibody was pooled together and treated with 40% saturated (∼4.32 M) Ammonium Sulfate to salt-out the conjugated antibodies from the unconjugated oligos still present in the antibody solution. After the majority of the remaining oligos were removed, the pooled solution was passed through a 50 kDa MWCO filter (EMD Millipore: UFC505024) 3-5 times. The complete protocol can be found at https://phospho-seq.com/post/protocol_1/PhosphoKit_Protocol.pdf.

### FlexPlex experimental workflow

In addition to the summary below, we have published a detailed protocol for FlexPlex here:https://phospho-seq.com/post/protocol_6/FlexPlexProtocol.pdf. We also include probe, hashtag oligo, and antibody-derived tag sequences in Supplementary Table 1.

For each condition in each experiment, two million cells are harvested from growth media resuspended in 450 μl of PBS and passed through a 40 μm strainer to remove clumped cells. Cells were fixed by adding 30 μl of 16% formaldehyde (Sigma: F8775-25ML) (final concentration, 1% FA in PBS). Cell suspensions were left to fix for 10 minutes at RT, with inversion every 3 minutes. The fixation reaction was quenched by adding 68.5 μl 1M glycine and filling the tubes with ice-cold PBS. The suspensions were centrifuged for 5 minutes at 400 x g at 4C, after which the supernatant was removed. The cells were resuspended in 1 ml PBS and the centrifugation was repeated. After the second centrifugation, the cells were resuspended in 100 μl of lysis buffer (10 mM Tris-HCl, 10 mM NaCl, 3.33 mM MgCl_2_, 0.1% NP-40 (Thermo: 28324), 1% BSA, 0.1 mM DTT (Invitrogen: y00147), 200U NxGen RNAse Inhibitor (Lucigen: 30281-1) in H_2_O and incubated on ice for 5 minutes for permeabilization. After 5 minutes, 1 ml of wash buffer #1 (10 mM Tris-HCl, 10 mM NaCl, 3.33 mM MgCl_2_, 1% BSA, 0.1 mM DTT, 100U NxGen RNAse Inhibitor in H_2_O) was added and cells were centrifuged for 5 minutes at 500 x g after which the supernatant was discarded and the cells were resuspended in 100 μl blocking buffer per hashing group and placed in a tube rotator at RT for 30 minutes. Blocking buffer consisted of 100 μg 30 nt blocking oligo, (NNNNNNNNNNNNNNNNNNNNNNNNNNNNN/3ddC/), 0.1 mM DTT, 100U NxGen RNAse Inhibitor and 3% BSA in PBS. For hashing, we spiked in 15 μl of 200 uM unique OligoHash per hash group. OligoHashing takes advantage of the innate electrochemical stickiness of the fixed cell intracellular environment, in which negatively charged DNA oligos will non-specifically bind to positively charged pockets within the cell ^26^. OligoHashes were designed to be capturable by the 10x Flex gel bead oligos by incorporating capture sequence 1 and uniquely amplifiable within the product library by incorporating a Nextera Read 2 PCR handle (Table S1). As a UMI for each molecule is added via the Flex chemistry, OligoHashes only need to have the hash index between the capture sequence and PCR handle, reducing the cost over other hashing methods by reducing the number of nucleotides needing to be synthesized. OligoHashes were ordered from IDT with the following sequence: CGGAGATGTGTATAAGAGACAGXXXXXXXXXXXXXXXGCTTTAAGGCCGGTCCTAGC*A*A where the Xs represent the unique OligoHash barcode.

While in the blocking and hashing step, the primary antibody pool was incubated with single-stranded DNA binding protein (Promega: M3011) to help reduce off-target effects ^14,23^. Briefly, 8 μg of SSB was added per 1 μg of antibody in a solution of 1x NEB buffer 4 (NEB:B7004S) for 30 mins at 37 C. After this incubation, BSA and PBS were added to make a final solution of 3% BSA in 1X PBS. Finally, 0.1 mM DTT and 100U/100 μl NxGen RNAse Inhibitor were added to the staining solution.

After blocking, cells were centrifuged at 600 x g for 5 minutes and washed with a wash buffer of 3% BSA and 0.1% Tween in PBS twice. After the second wash, the individually hashed pools were combined in wash buffer and centrifuged again before resuspending in the previously prepared primary antibody staining solution with a maximum of two million cells. The cells were placed on a tube rotator at room temperature for 1 hour for primary antibody staining.

After 1 hour, the cells were centrifuged and washed three times in the same wash solution from the blocking step. After the final supernatant was removed, the cells were input into the manufacturer recommended 10x Flex workflow starting with 4% formaldehyde fixation, which acts as a second fixation step.

For each experiment, we used 10x Flex v1 kit for singleplexed samples with feature barcoding. This kit enables the capture of both RNA probes and TotalSeq-B conjugated antibodies within the same cells. We made minor modifications to the protocol to facilitate the capture of guides and OligoHashes. First, because we used OligoHashing in each of our experiments, which enables high throughput doublet demultiplexing ^25^, we can superload the microfluidic chip over the recommended loading quantity. Next, we spike in 1 μl of 0.2 μM an OligoHash additive primer into the pre-amplification step, in order to preamplify the OligoHashes alongside the mRNA, guides and ADTs. The sequence for that primer is: CGGAGATGTGTATAAGAGAC*A*G. Once the preamplification is finished, the final products are eluted into 100 μl of elution buffer. Each modality will use 20 μl of this solution as the basis for modality specific PCR. Transcriptome and ADTs are amplified according to the protocol as written. OligoHashes are amplified by using a Nextera Read 2 index primer (CAAGCAGAAGACGGCATACGAGATNNNNNNNNGTCTCGTGGGCTCGGAGATGTGTATAA G) and SI-PCR primer (10 μl of 10 μM each). Each modality is amplified under the same PCR conditions as written for the transcriptome, with 12-14 cycles for ADTs and OligoHashes. For guides, their sparsity requires a second preamplification step. This PCR uses 1 μl each of 10 μM TruSeq Read1 primer (CTACACGACGCTCTTCCGATCT) and TruSeq-GDO Read2 primer (GTGACTGGAGTTCAGACGTGTGCTCTTCCGATCTGCTATGCTGT) and 20 μl of the preamplification elute for 8 cycles. Afterwards, products are eluted into 30 μl following a 1.8x SPRI cleanup. After elution, 7 further cycles of PCR are performed using indexing primers. Sequencing for all libraries was performed using a NextSeq 550 instrument with the read structure of 28/8/50. Optimal read numbers per cell vary, but are generally 10,000 for gene expression, 5,000 for ADT, 2,000 for OligoHash and 2,000 for guides. Guide RNA sequences for all experiments are listed in Table S4.

### FlexPlex computational processing

Transcriptomic data was aligned and quantified using CellRanger 8.0 to generate count matrices. These matrices were imported into Seurat v5.2.1 ^60,61^ for further processing including log-normalization and dimensionality reduction using default parameters. Guides (GDOs), OligoHashes (HTOs) and ADTs were quantified using a custom script that scanned the FASTQ files for cell barcodes, unique molecular identifiers and known index sequences, building a counts matrix from this information. Each version of the quantification script can be found at https://phospho-seq.com/files/quant.ipynb.

HTOs were normalized using CLR normalization, and we performed demultiplexing and doublet detection using the MULTISeqdemux function with default parameters. Only cells classified as singlets were kept. GDOs were normalized using CLR normalization, and we performed demultiplexing and doublet detection using the MULTISeqdemux function with quantile = 0.5. Only cells classified as singlets for the gene level counts were kept. Finally, ADTs were normalized using CLR normalization. For the pilot dataset, we additionally removed other cells multiplexed in for testing other experimental parameters, leaving the total number of processed cells at 1,665. We used the same workflow for the ‘glossary’ and ‘validation’ experiments.

### Genome-wide Pooled Screen

K562 cells were cultured as described above and expanded to 180 million cells. Lentivirus containing genome-wide targeting libraries, with six guides per gene, was applied to the cells with a target multiplicity of infection of 0.3. Puromycin was added to the cell culture one day after infection and selection continued for nine days, with culture expansion and media changes every three days. On the ninth day, dead cells were removed from the culture with a dead cell removal kit and columns (Miltenyi Biotec: 130-090-101;130-042-401) and live cells were returned to culture to recover.

On the day of the experiment, cells were processed similarly through the primary antibody staining to a FlexPlex experiment with minor modifications. First, all volumes were scaled up 50-fold for fixation, permeabilization and staining. Further, all RNAse inhibitor and DTT were left out of each buffer as RNA quality is not integral to success of the assay. Blocking was performed in 3% BSA in PBS, with no additional additives. Finally, primary antibody staining was performed using a 1:250 dilution of unconjugated primary pRPS6 antibody (Biolegend: 608602) for 1 hr on a tube rotator at room temperature. After primary staining, cells were washed twice in 3% BSA + 0.1% Tween in PBS and stained with FITC conjugated secondary antibody (Biolegend: 406605) for 30 minutes on a tube rotator at room temperature, protected from light. Cells were washed three times and resuspended in MACS buffer (0.5% BSA + 2mM EDTA in PBS) for sorting.

All sorting was performed on a Sony SH800 cell sorter. Cells were sorted simultaneously according to gates representing the top and bottom 10% of pRPS6 signal across all cells. The number of cells targeted to sort per bin was 5.7 million, which represents 500 cells per guide. After sorting, the genomic DNA from cells was collected using the Qiagen DNeasy Blood and Tissue DNA extraction kit (69504) using manufacturer’s instructions with a minor modification to account for the cell fixation. Cells were centrifuged and resuspended in 180 ul buffer ATL and 20 μl proteinase K and incubated overnight at 56C to ensure complete nucleosome digestion. The remainder of the protocol was performed as written in manufacturers instructions.

Guides were amplified from genomic DNA with the following PCR mix: Up to 10 μg of genomic DNA, 10 μl reaction buffer, 8 μl dNTPs, 1.5 μl ExTaq (all from Takara: RR001A), 5 μl 10 uM P5 Primer pool (AATGATACGGCGACCACCGAGATCTACACTCTTTCCCTACACGACGCTCTTCCGATCTyXX XXXXXXtcttgtggaaaggacgaaacaccg), 5 μl 10 uM P7 primer (CAAGCAGAAGACGGCATACGAGATXXXXXXXXGTGACTGGAGTTCAGACGTGTGCTCTTC CGATCTyCCGACTCGGTGCCACTTTTTCAA) and water up to 100 μl. The ‘y’ in each primer represents a stagger base that can be expanded up to six bases that is necessary for sequence library complexity. The ‘X’ in each primer represents the 8 base index used for sequence demultiplexing. The PCR conditions are 95C for 1 minute, 95C for 30 seconds, 53C for 30 seconds, 72C for 30 seconds, repeating steps 2-4 for 28 cycles, with a final extension of 72C for 10 mins. PCR products were cleaned up with a double sided SPRI of 0.6x and 1.6x to remove primers and remaining genomic DNA. Libraries were quantified using a qubit and run on a NextSeq 550 instrument with a Read1: 60/Index1: 8 format.

Sequencing data was demultiplexed, reads were trimmed using Cutadapt v4.4 ^62^ and guides were quantified and analyzed using MAGeCK v0.5.9.5 ^30^, which includes statistical analysis and Robust Rank Aggregation scores.

### Building and Applying Predictive Models of pRPS6 from scRNA-seq data

To build predictive models, log-normalized counts matrices from the glossary dataset (subset for genes present in the genome-wide perturb-seq dataset) and CLR-normalized pRPS6 values were input into the cv.glmnet function from the glmnet package (v4.1) in R ^63^ with the following parameters: alpha = 0 (Ridge) ^36^ or 1 (LASSO) ^35^, family = “gaussian”, type.measure = “class”, nfolds = 10. The lambda with the minimum cross-validation error (lambda.min) was extracted from this model and the final model was determined with the same inputs and the following parameters: alpha = 0, family = “gaussian”, lambda = lambda.min.

To determine the most important components of the model, we extracted the coefficients from the model, using the ‘coef’ function and s = lambda.min as input. We multiplied these coefficients by average log-counts for each gene across the prediction dataset. This product reflects the amount that each gene, at its average expression level, contributes to the pRPS6 prediction. We refer to this as the ‘model contribution score’. To apply the predictive models to new datasets, we extracted log-normalized count matrices, filtered them for model genes, and used the Base R ‘predict’ function to generate pRPS6 predictions for each cell. To determine perturbations that produce significant predicted changes in pRPS6 we applied a t-test between the “non-targeting” cells and each perturbation, correcting for multiple testing with the Benjamini-Hochberg method.

To determine the perturbations that would go into the validation dataset, we first filtered the 586 significant perturbations for predicted pRPS6, selecting the top 200 downregulating and 50 upregulating by log-fold change. Conscious that the strongest perturbations would produce similar transcriptional results, we opted to leverage the perturbation-level clustering from GWPS ^20^ and our own clustering for manual curation. This produced a selection of 100 perturbations with diverse transcriptomic consequences, to which we added 20 additional perturbations as positive controls previously linked to mTOR signaling even if they were not hits in our ‘in silico’ screen (Table S3).

### Western Blotting

K562 cells were harvested in lysis buffer containing 2 mM EDTA (Sigma: E5134), 2 mM EGTA (Sigma: E3889), 1% Triton-X (Sigma: T8787), and 0.5% SDS (Sigma: 71736) in 1x PBS with EZ Block phosphatase inhibitor cocktail (BioVision: K273-1) and Complete mini EDTA-free protease inhibitor cocktail (Sigma: 4693159001). Total protein was determined by BCA assay (ThermoFisher: 23227) and 15 μg of protein in Nupage LDS sample buffer (Thermo: NP0007) were loaded onto Novex 4-12% Tris-Glycine minigels (Thermo: XP04125). Proteins were transferred to PVDF membranes (EMD Millipore: IPFL07810), blocked in 5% milk in TBS-Tween for one hour at room temperature (RT), and incubated with primary antibodies diluted in 5% milk in TBS-Tween overnight at 4°C. The following day, membranes were incubated with HRP-conjugated secondary antibodies (Jackson ImmunoResearch: 111-035-003;115-035-003) for one hour at RT, washed, incubated with chemiluminesence substrate (Thermo: 34577) and scanned on a Licor C-digit device. The concentration and catalog number of primary antibodies were as follows: ATF4 1:1000 (Cell Signaling: 11815), Histone H3 1:2000 (Cell Signaling: 4499), pRPS6 1:1000 (BioLegend: 608602), RPS6 1:1000 (BioLegend: 691802).

## Data availability

Seurat is freely available as open-source software packages at: https://github.com/satijalab/seurat

FlexPlex datasets for the pilot, glossary, and validation experiments that were generated for this manuscript are available at: https://doi.org/10.5281/zenodo.16763127

## Author Contributions

J.D.B. and R.S. conceived of the study and wrote the manuscript. All experiments were performed by J.D.B, A.B. and C.D. J.D.B performed computational analysis with guidance and supervision from I.N.G. and R.S.

## Acknowledgements

The authors would like to thank all the members of the Satija Lab for thoughtful discussions related to this work. J.D.B. is a postdoctoral fellow of the Jane Coffin Childs Memorial Fund for Medical Research. I.N.G. is the Kenneth G. Langone Quantitative Biology Fellow of the Damon Runyon foundation (DRQ-21-24). This work was supported by the Chan Zuckerberg Initiative (EOSS-0000000082, HCA-A-1704-01895 to R.S.), and the NIH (RM1HG011014-02, 1OT2OD026673-01, R01HD096770, R35NS097404 to R.S).

---

**Figure S1:**
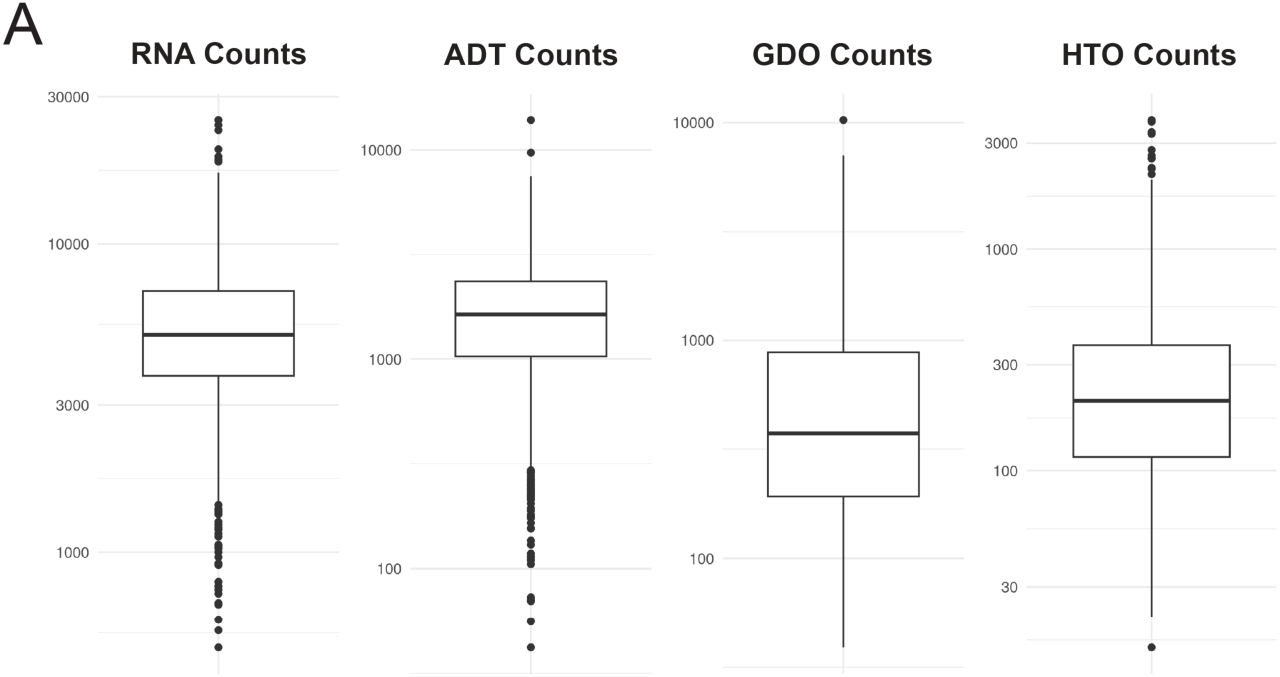
**a)** Box plots of RNA, ADT, GDO and HTO counts per cell in the FlexPlex pilot experiment.

**Figure S2:**
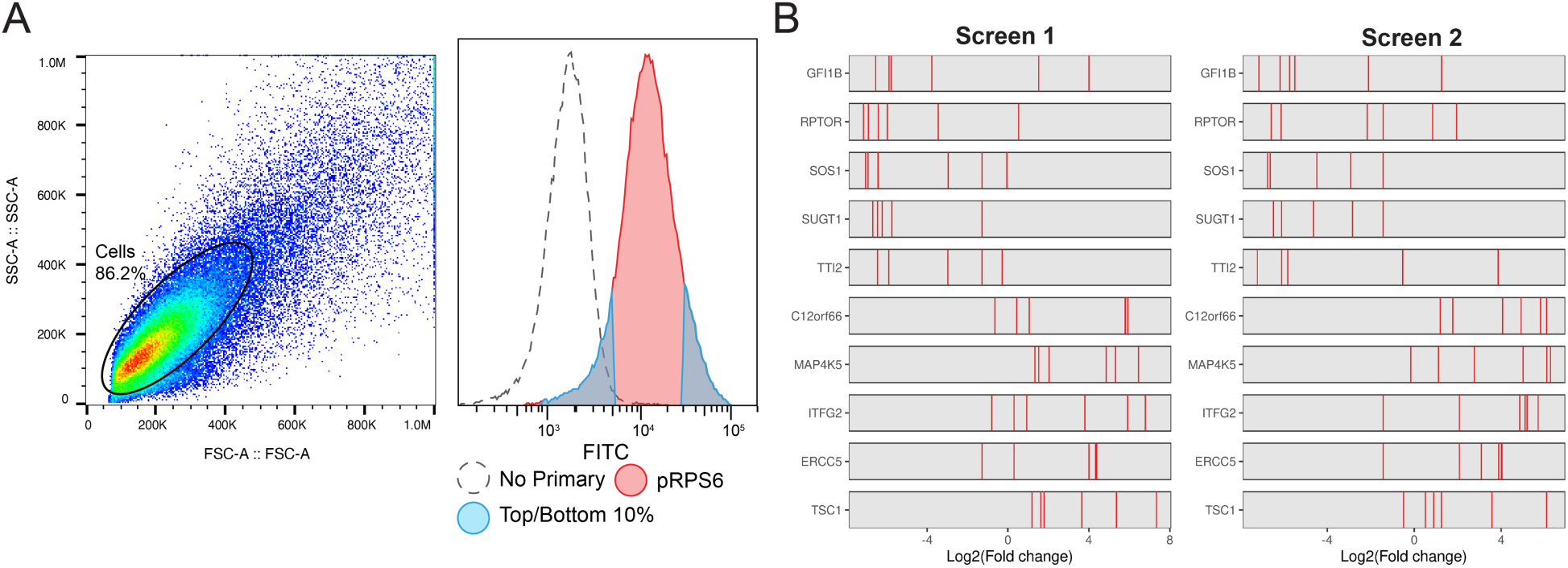
**a)** Sorting gates of genome-wide bulk screen sorting on pRPS6 showing sorted fractions and unstained control. **b)** GuideRank Plots showing top hits reproducing across two replicates. Each red line represents the fold-change of an individual gRNA, grouped by target gene.

**Figure S3:**
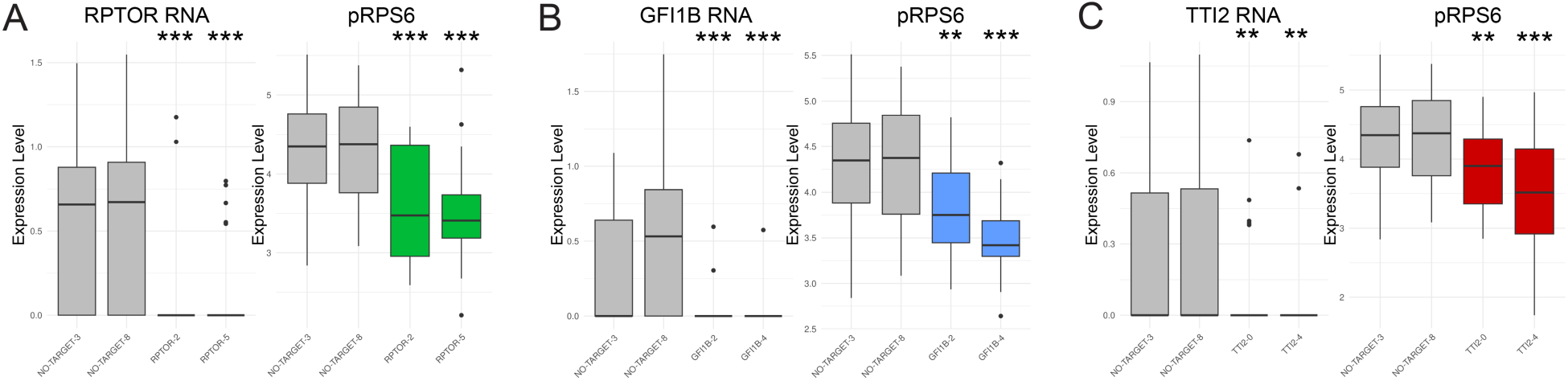
**a)** Boxplots showing guide reproducibility for the target RNA and pRPS6 between two non-targeting and two guides for **(a)** RPTOR, **(b)** GFI1B and **(c)** TTI2. All statistical tests are two-sided t-tests adjusted using Benjamini-Hochberg correction for multiple comparisons. Significant differences between compared to non-targeting controls indicated by *** p <0.001,**p<0.01,*p<0.05. For all boxplots, the upper and lower bounds represent 75th and 25th percentiles respectively and the center line represents the median.

**Figure S4:**
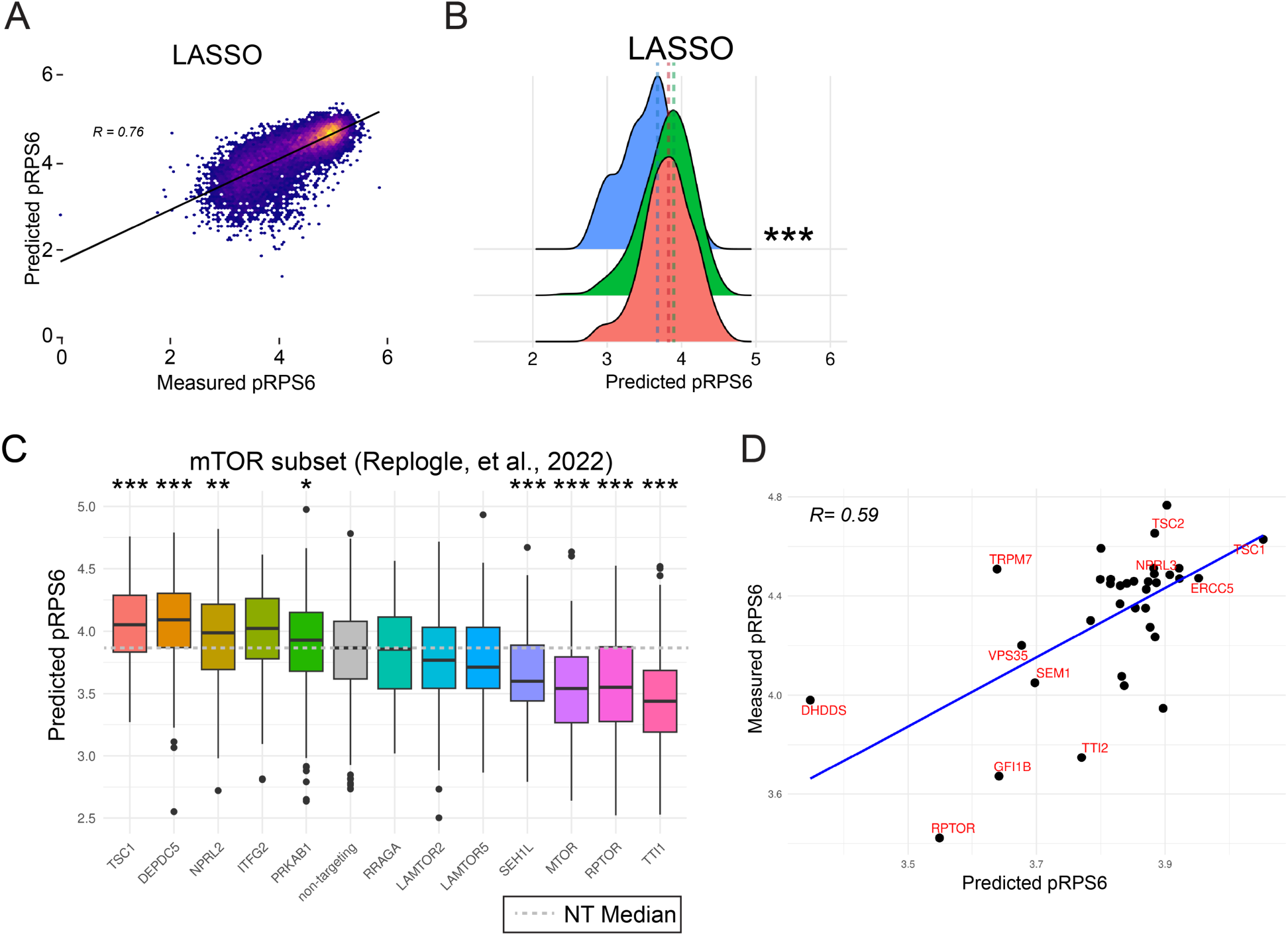
**a)** Correlation density plot comparing predicted pRPS6 to measured pRPS6 on a validation dataset for the LASSO modeling method (R=0.76, p < 0.0001). **b)** Density plot demonstrating the predicted pRPS6 differences between RPTOR, TSC1 and non-targeting perturbations across three linear modeling methods using transcriptional data from genome-wide perturb-seq (results for Ridge and OLS are shown in Fig. 4e). **c)** Boxplot of predicted pRPS6 across a subset of mTOR-related perturbations in genome-wide perturb-seq. **d)** Correlation scatter plot of median predicted pRPS6 values from GWPS and measured pRPS6 values from the glossary (R=0.59; p = 0.0001). All statistical tests are two-sided t-tests adjusted using Benjamini-Hochberg correction for multiple comparisons. Significant differences between compared to non-targeting controls indicated by *** p <0.001,**p<0.01,*p<0.05. For all boxplots, the upper and lower bounds represent 75th and 25th percentiles respectively and the center line represents the median.

**Figure S5:**
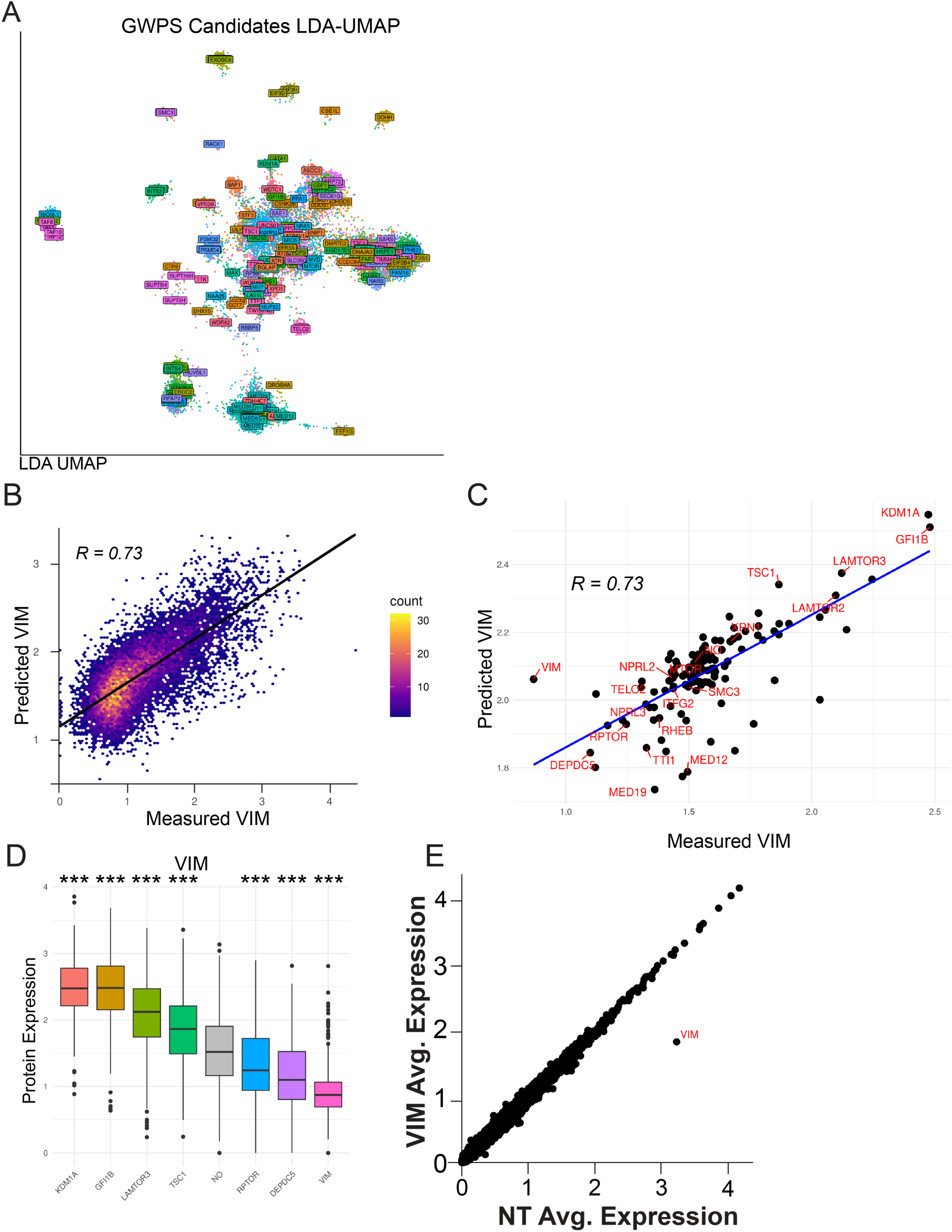
**a)** LDA UMAP of cells from 250 perturbations predicted to regulate pRPS6 from GWPS. We clustered this space which, along with the clustering from the original manuscript, was used to select a diverse set of 120 experimental perturbations for follow-up. **b)** Correlation density plot comparing predicted Vimentin levels to measured Vimentin levels on a validation dataset (R=0.73, p < 0.0001). **c)** Correlation of the medians of predicted vs. measured Vimentin levels for each perturbation in the validation experiment (R= 0.73, p < 0.0001). **d)** Boxplot of VIM protein levels for a subset of perturbations in the validation experiment. **e)** Scatter plot of average gene expression for each gene between non-targeting cells and VIM perturbed cells. All statistical tests are two-sided t-tests adjusted using Benjamini-Hochberg correction for multiple comparisons. Significant differences between compared to non-targeting controls indicated by *** p <0.001,**p<0.01,*p<0.05. For all boxplots, the upper and lower bounds represent 75th and 25th percentiles respectively and the center line represents the median.

**Figure S6:**
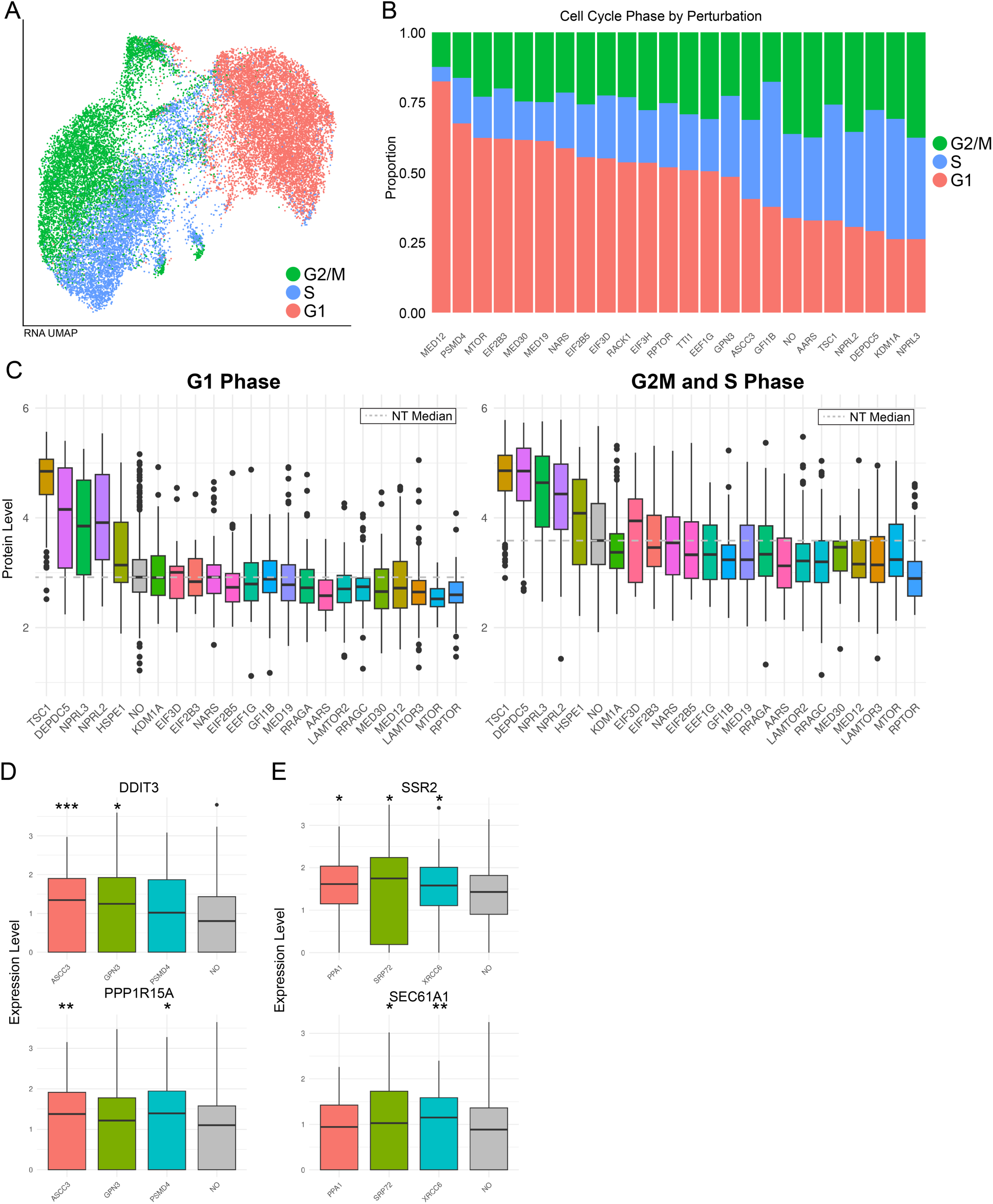
**a)** RNA UMAP of cells across validation experiment grouped by cell cycle phase. **b)** Proportional bar plot of cell cycle phase across a subset of perturbations. **c)** Boxplot of measured pRPS6 among a subset of perturbations, after splitting into proliferating (G2M+S) and G1 phase groups. **d)** Boxplot of general stress related transcripts across growth-related perturbations. **e)** Boxplot of IREα/ATF6 stress related transcripts across UPR-related perturbations. All statistical tests are two-sided t-tests adjusted using Benjamini-Hochberg correction for multiple comparisons. Significant differences between compared to non-targeting controls indicated by *** p <0.001,**p<0.01,*p<0.05. For all boxplots, the upper and lower bounds represent 75th and 25th percentiles respectively and the center line represents the median.

